# AlterNet: Alternative splicing-aware gene regulatory network inference

**DOI:** 10.1101/2025.11.21.689745

**Authors:** Juliane Hoffmann, Julia Wallnig, Ziheng Dai, Olga Tsoy, David B. Blumenthal, Anne Hartebrodt

## Abstract

Gene regulatory networks (GRNs) help decode biological systems by identifying how genes interact and regulate cellular processes. However, conventional GRN inference methods operate at the gene-level, overlooking transcript-level variability introduced by alternative splicing (AS). In this work, we present AlterNet, the first GRN inference and annotation pipeline which considers isoform of transcription factors as distinct regulators in a GRN. AlterNet builds on the GRNBoost2 inference algorithm, and includes a transcript plausibility and annotation workflow. We applied AlterNet to expression data from heart tissue, including samples from donors with normal heart function and from patients with different types of cardiomyopathy. The resulting isoform-level GRNs uncovered highly relevant regulatory interactions not detectable at the gene-level. Overall, AlterNet infers transcript-level regulatory network which enable the discovery of novel, biologically relevant regulatory interactions that remain hidden in gene-level GRNs. The source code of AlterNet is available on GitHub https://github.com/bionetslab/AlterNet, and can be installed as a Python package.

## 1 Introduction

Alternative splicing (AS) is a process during gene expression that allows a single gene to produce multiple proteins (isoform). AS takes place during or after [20] and involves the process of including or excluding specific exons from the final RNA sequence [1]. Through these alterations of the original RNA sequence, different transcript variants of the same genes are obtained, which encode different protein isoforms that can be functionally diverse. Protein isoforms from the same gene cannot only have different interaction partners [14], and may also assume a different functional role in gene regulation [12]. In particular, AS can lead to differential behaviour transcription factor (TF) isoforms, even if the sequence variations occur outside of annotated functional domains [12].

TFs are regulatory proteins that bind to specific DNA sequences and thereby modulate the expression of target genes in the vicinity of their binding sites [20]. TF-mediated transcriptional regulation can be represented by gene regulatory networks (GRNs). In a GRN, nodes represent genes (TFs and target genes) and directed edges from TFs to their target genes represent regulatory relationships. A variety of algorithms are available to infer GRNs from gene expression data [16]. Existing GRN inference methods use diverse techniques such as tree-based machine learning [8,18], linear regression models [21], ordinary differential equations [15], or mutual information [11], but all share one common limitation: They model GRNs at gene level and neglect the complexity introduced by AS. The gene-level approach of existing GRN inference tools is oblivious to potentially important differences between TF isoforms, such as the TF isoform-specific regulation patterns identified by a recent study on in GTEx tissue data [12]. Furthermore, there are existing case studies, which consider alternative splicing in the study of gene expression at network level [30] which indicate the general interest of this topic.

To address this limitation of existing approaches, we developed AlterNet — the first AS-aware GRN inference method which allows to identify regulatory links involving TF isoforms that remain invisible in conventional gene-level GRN inference. AlterNet consists of four steps: In the first step, TF-target gene links are inferred with a variant of the popular GRNBoost2 [18] method, in parallel for transcript- and gene-level TF representations (target genes are always represented at gene level because target gene isoforms are controlled by splice factors that act downstream of TF-mediated transcriptional regulation). In the second step, AlterNet categorizes the the inferred transcript- and gene-level edges into diferent groups, which are then filtered in the third step to restrict to *prima facie* plausible interactions. In the fourth step, the plausible interactions are augmented with isoform function annotations from the APPRIS [25] and DIGGER [14] databases to facilitate the identification and prioritization of biologically plausible regulatory connections and isoforms with unique features.

We validated AlterNet on gene expression data from human heart tissue, motivated by the fact that AS has been found to be a relevant mechanism in cardiomyopathies [3,26]. Cardiomyopathies affect the stucture and function of the heart. Dilated cardiomyopathy (DCM) is the most prevalent form and is characterized by a dilation of the left or both ventricles [3]. Hypertrophic cardiomyopathy (HCM) is the second most common form with a thickening of the ventricular wall. Here, we use a gene expression dataset from the MAGNet consortium [5], with 166 DCM and 28 HCM samples harvested during surgery and 166 samples from non-failing donor hearts (NF). An analysis of the domains contained in TF isoforms that are predicted as regulators by AlterNet showed an enrichment in domains mediating DNA-protein interaction, lending evidence to the hypothesis that these isoforms act as TFs that regulate transcription. Moreover, gene set enrichment analysis on the target genes in, respectively, AlterNet’s isoform-specific GRNs and canonical gene-level GRNs computed with GRNBoost2 showed a shift from generic terms obtained for the canonical GRNs to more condition-specific cardiomyopathy-related terms in the isoform-specific GRNs.

## 2 Results

### 2.1 Overview of the AlterNet pipeline

Figure 1 provides an overview of the AlterNet pipeline, which consists of four steps: first, the inference of a canonical GRN and an AS-aware GRN, second, an edge categorization step, third, a plausibility filtering step which aims to prioritize statistically relevant edges, and fourth, an annotation workflow to contextualize isoforms that act as regulators in the filtered network. Input to the first step is a transcript expression matrix *X*^ℐ^ ∈ ℝ^*n×*|ℐ|^ ( is the set of isoforms, *n* the number of samples), an isoform-to-gene mapping *σ* :, ℐ → 𝒢 and a set ℛ ⊆ 𝒢 of TFs that serve as candidate regulators. Using *σ*, AlterNet aggregates *X*^ℐ^ to a gene expression matrix *X*^𝒢^ ∈ ℝ^*n×*|𝒢|^ and then calls GRNBoost2 in two mode: The first mode takes *X*^𝒢^ and ℛ as input and yields a canonical GRN *G*^*c*^ with edges (*r, g*) from TFs *r* ∈ ℛ to target genes *g* ∈ 𝒢. The second mode takes *X*^ℐ^, *X*^𝒢^, and ℛ^ℐ^ = *σ*^−1^[ℛ] as input and returns an AS-aware GRN *G*^*a*^ with edges (*i, g*) from TF isoforms *i* ∈ ℛ ^ℐ^ to target genes *g* ∈ 𝒢. Note that, since AS acts downstream of transcriptional regulation, targets are modeled as genes and not as isoforms also in *G*^*a*^.

**Fig. 1.**
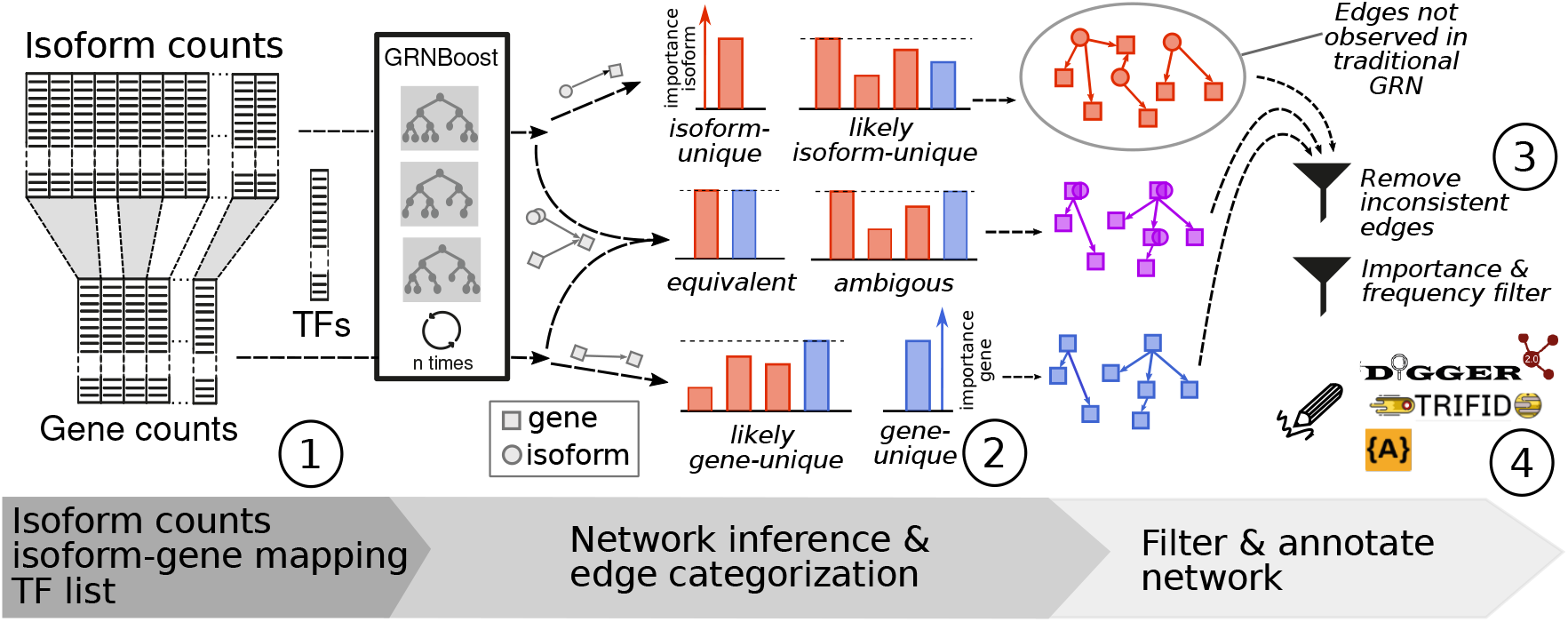
General workflow of AlterNet. AlterNet takes transcript expression data and a list of TFs as input for inference of a gene-level canonical GRN and an isoform-level AS-aware GRN, using a customized version of the algorithm GRNBoost2 [18]. The inference is repeated a configured number of times, then the edges are aggregated to one gene-level and one isoform-level consensus network. The inferred network edges are categorized into the categories *isoform-unique, gene-unique* and *common* interactions, and common interactions are further subdivided into *likely isoform-unique, likely gene-unique, equivalent*, and *ambiguous* interactions. Several filtering steps are applied to remove inconsistent or implausible edges from the different categories, and the remaining plausible interactions are annotated with information on functional relevance as well as exon and domain usage obtained from the DIGGER [14] and APPRIS [25] databases.

In the second step, we classify edges in the two networks as *isoform-unique, gene-unique*, and *common*. Edges (*i, g*) and (*σ*(*i*), *g*) are labeled as common if (*i, g*) appears in the AS-aware GRN and (*σ*(*i*), *g*) in the canonical GRN. The remaining edges are labeled as isoform-unique if they only appear in the AS-aware GRN and as gene-unique if they only appear in the canonical GRN. Based on the distributions of inferred edge importance weights in the canonical and AS-aware GRNs, the common edges are further subdivided into *likely isoform-unique, equivalent, ambiguous*, and *likely gene-unique* edges (Figure 1, middle). Of those, the likely isoform-unique edges are particularly relevant, as they constitute regulatory links that are stronger in the AS-aware GRN than in the canonical GRN and are thus easily overlooked when focussing on the edges with the highest weights in the canonical GRN. Edges that are either isoform-unique or likely isoform-unique (i. e., the set of all links that tend to remain invisible in gene-level GRN inference) are called *isoform-specific*.

In the third step, we filter the edges from the different categories via a *frequency filter*, an *equivalence filter*, a *dominance filter*, a *foldchange filter*, and an *importance filter* to retain a small set of robustly detectable and statistically plausible edges. The frequency filter removes all edges of all categories that are not robustly detectable via several calls to GRNBoost2. The equivalence and dominance filters remove all isoform-unique and gene-unique edges where the TF maps to a single isoform or to a dominant isoform that accounts for almost all of the transcript counts. In these cases, the TF-level and TF isoform-level expression vectors are identical (equivalence filter) or close to identical (dominance filter), and corresponding regulatory links that appear either only in the AS-aware GRN or only in the canonical GRN are thus considered false positives. Using an analogous reasoning, the foldchange filter removes common equivalent edges (i. e., edges found in both the canonical and the AS-aware GRN where TFs map to a single isoform) for which the inferred importance weights in the AS-aware and in the canonical GRN differ substantially. Finally, the importance filter sorts all edges retained in the previous filtering steps by the importance weights inferred by GRNBoost2 and then only keeps the edges at top of the list. The number of GRNboost2 calls, and the percentage of top edges is user defined (here we choose 10 calls, and use the top 20% edges).

In the final step, the isoforms involved in the filtered GRNs are annotated with domain and exon usage labels from DIGGER and functionally defined isoform categories *principal, alternative*, and *minor* from AP-PRIS. Moreover, we retrieve TRIFID scores [23] from APPRIS, which range between 0 and 1 and constitute machine learning-based estimates of the functional importance of isoforms. These functional annotations allow users to prioritize interactions that involve isoforms with specific functional characteristics such as unique exons or domains. Detailed explanations of all steps of the AlterNet pipeline are provided in Methods.

### 2.2 AlterNet yields concise networks of isoform-specific regulatory edges

Figure 2 shows the number of edges before and after filtering for the DCM, HCM, and NF datasets (datasets are represented by different shades of gray, blue, and red). The unfiltered AS-aware and canonical GRNs each contain 10s of millions of edges (Figure 2A) of which most are categorized as common edges (Figure 2B). The largest numbers of edges are removed by the frequency filter (compare the counts shown in Figure 2B and in the first rows in Figure 2C, E). The gene-unique edges are more likely to be false positives than the isoform-unique edges, as the equivalence filter removes almost half of the gene-unique but almost none of the isoform-unique edges (Figure 2C, E). Within the subset of common edges, the largest number of edges are gene-unique and equivalent, followed by ambiguous edges and likely isoform-unique edges (Figure 2D). Overall, we observe that the frequency and the importance filters remove the largest number of edges and that the equivalence and dominance filters are more relevant for the gene-unique than for the isoform-unique edges. After all filtering steps, the isoform-specific subnetworks for all three datasets contain fewer than 10000 high-confidence edges (Figure 2F).

**Fig. 2.**
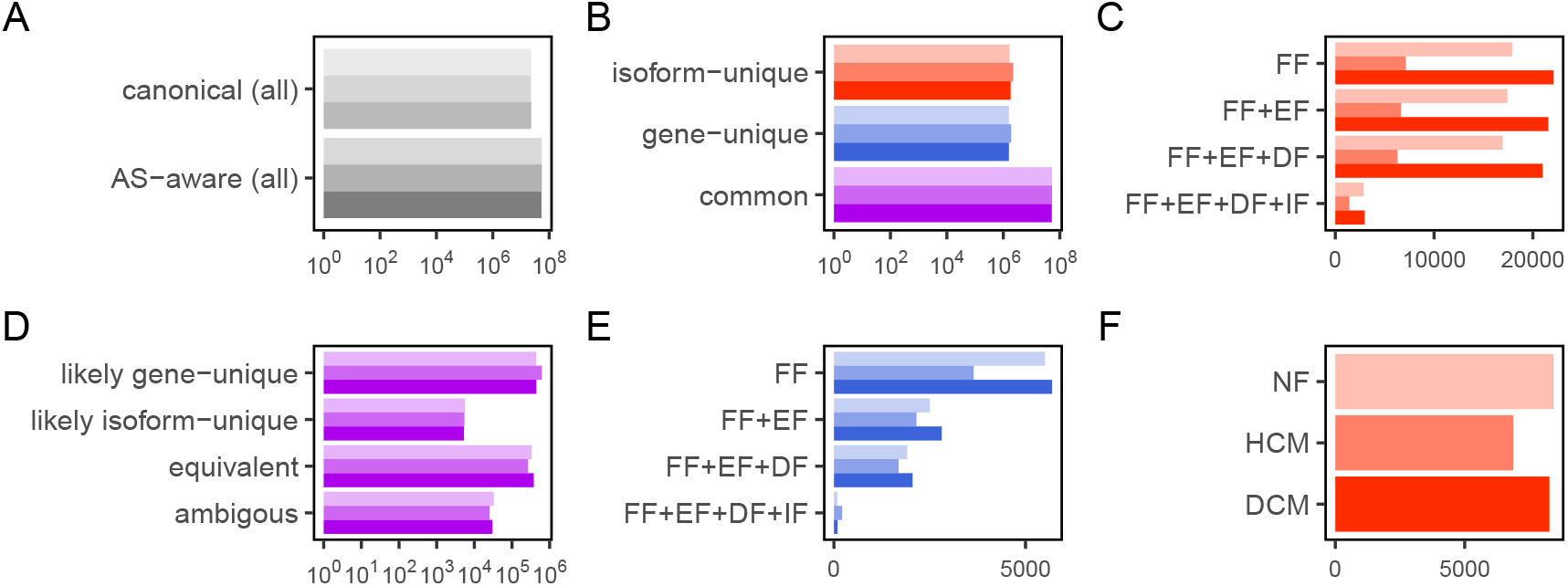
Effect of edge filtering for the DCM (dark), HCM (medium), and NF (light) datasets. (A) Total numbers of edges in the unfiltered canonical and AS-aware networks. (B) Numbers of edges in the disjoint sets of isoform-unique, gene-unique, and common edges. (C, E) the effects of different filters (FF: frequency filter, EF: equivalence filter, DF: dominance filter, IF: importance filter) on the isoform-unqiue (C) and gene-unique (E) edge sets. (D) Number of edges in each subcategory of the common edges after filtering (filtering steps not shown). (F) Numbers of isoform-specific edges in the three cohorts after applying all filtering steps.

### 2.3 TF isoforms involved in isoform-specific edges are functionally plausible

To assess if the TF isoforms involved in the filtered isoform-specific edges uncovered by AlterNet are functionally plausible, we carried out functional enrichment analysis to identify Gene Ontology (GO) [2,6] terms that are overrepresented in Pfam domains that are exclusively associated with these TF isoforms in comparison to domains associated with all of the TF isoforms appearing in the unfiltered AS-aware GRNs. The results for the three datasets are shown in Table 1. After Bonferroni correction, we found significant enrichment indicating DNA binding, zinc-ion binding, and vesicular transport-related functions. In particular, DNA binding indicates that the TF isoforms identified as source nodes in the isoform-specific subnetworks are indeed likely to be acting as transcriptional regulators.

**Table 1.** GO terms overrepresented in domains that are found exclusively in TF isoforms involved in isoform-specific interactions returned by AlterNet for the DCM, HCM, and NF datasets. *P*-values were computed with the one-sided hypergeometric test and adjusted for multiple testing via Bonferroni correction.

| GO term | Adjusted $P$-value | | |
| --- | --- | --- | --- |
|  | DCM | HCM | NF |
| DNA binding | 0.015 | 0.033 | $\geq 0.05$ |
| Golgi membrane | 0.013 | 0.018 | 0.0004 |
| RNA binding | 0.046 | $\geq 0.05$ | 0.003 |
| intra-Golgi vesicle-mediated transport | 0.013 | 0.018 | 0.0004 |
| protein modification process | 0.013 | 0.018 | 0.01 |
| zinc ion binding | 0.013 | 0.039 | $\geq 0.05$ |

To further evaluate the isoforme-specific interactions identified by AlterNet, we next analyzed the distributions of functional DIGGER and APPRIS annotations of the isoforms acting as regulators in the filtered isoform-specific subnetworks in comparison to isoforms that appear in the AS-aware GRNs that were filtered only with the importance filter and thus also contain TF isoform-target links corresponding to TF-target links that are uncovered by conventional gene-level GRN inference. Figure 3 shows the results for the DCM datasets, the corresponding results for the HCM and NF datasets are shown in Figure A1 and A2 in the appendix. Most of the isoforms in the filtered isoform-specific network have a unique exon, but only few have unique domains (Figure 3A, B). The APPRIS annotations show a slight shift from principal to minor isoforms and from higher towards lower TRIFID scores in the isoform-specific network (Figure 3C, D). This is an expected result, as it implies that isoforms that are estimated to be functionally important based on existing knowledgebases and machine learning models trained on them are often already covered by genelevel GRNs. To assess the plausibility of isoforms uncovered by AlterNet which, according to APPRIS, have lower functional evidence, we therefore focused on the non-principal isoforms in the filtered isoform-specific subnetworks and asked whether more of them have unique or missing Pfam domains with respect to the corresponding principal isoforms than expected by chance. In comparison to background distributions that were computed by randomly sampling size-matched subnetworks from the filtered AS-aware GRN, we observe enrichments of both isoforms with unique and of isoforms with missing domains (Figure 3E, F). These results lend further evidence to the hypothesis that TF isoforms involved in isoform-specific interactions uncovered by AlterNet indeed have distinct functional profiles that could explain their involvement in regulatory links that are missed when analyzing transcriptional regulation at gene level only.

**Fig. 3.**
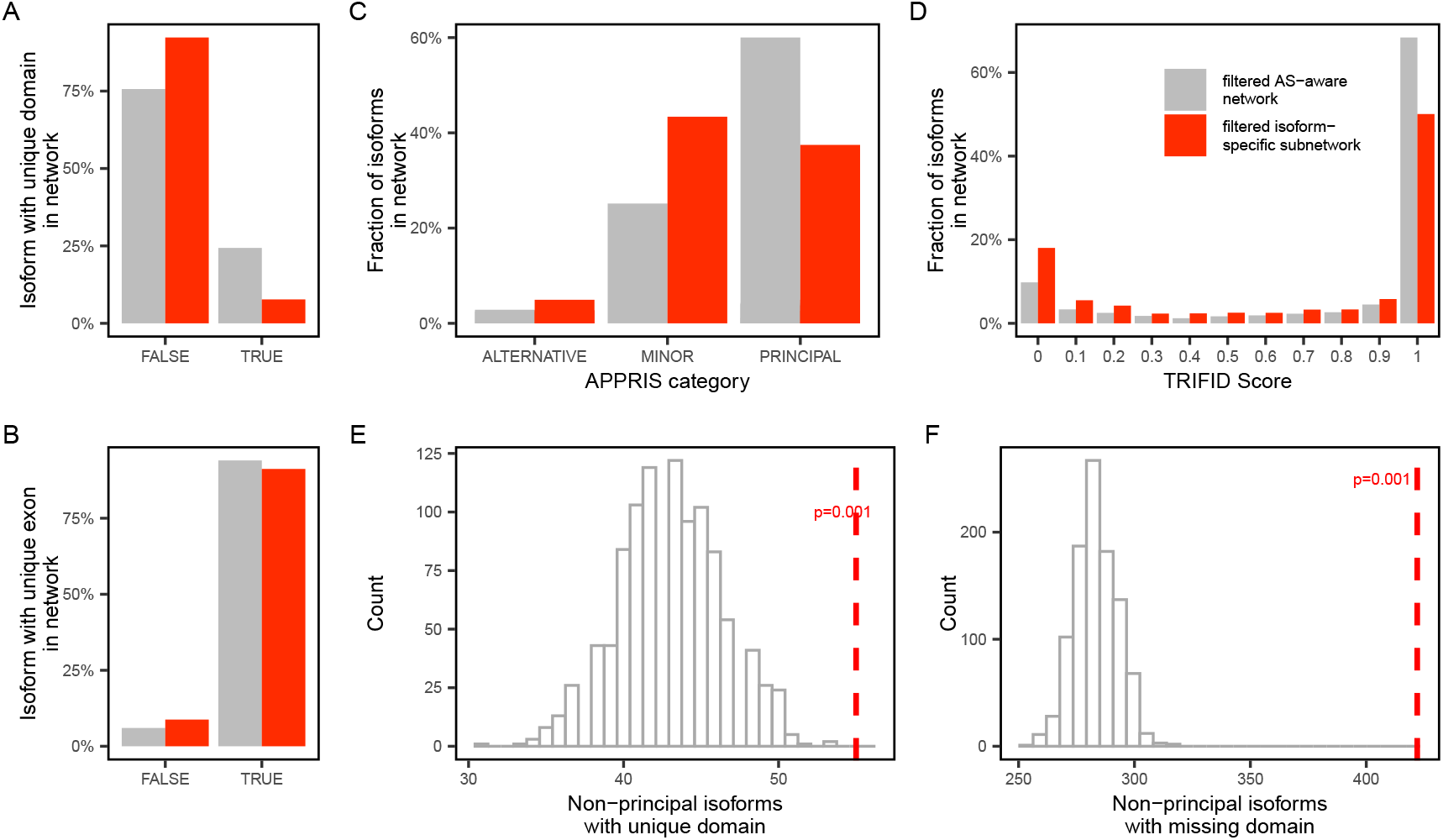
Analysis of TF isoforms in AlterNet’s filtered isoform-specific network obtained for the DCM dataset in comparison to the filtered AS-aware network. (A, B) Numbers of isoforms with unique exons and unique domains. (C, D) Distributions of the APPRIS isoform categories TRIFID scores. (E, F) Numbers of non-principal isoforms in the isoform-specific network with unique domains (E) and missing domains (F) against background distributions generated by randomly sampling an identical number of edges from the filtered AS-aware GRN. Empirical *P*-values were computed as explained in Section 4.3.

### 2.4 Targets of isoform-specific edges are more condition-specific than targets of gene-level regulatory edges inferred by vanilla GRNBoost2

To directly test AlterNet against classical gene-level GRN inference with GRNBoost2, we next compared the top 100 targets in AlterNet’s filtered isoform-specific networks (recall that these networks contain edges that are typically missed by classical GRN inference) to the top 100 targets in canonical GRNs inferred by vanilla GRNBoost2, using gene set enrichment analysis with g:Profiler [10]. The targets were ranked by their in-degree weighted by median importance. The results for the DCM dataset are shown in Table 2, corresponding results for the HCM and NF datasets are contained in Table A1 and A2 in the appendix. Among the significant highlighted terms, AlterNet retrieves several terms directly related to cardiac development (“cardiocyte differentiation”, “atrioventricular node cell fate commitment”), as well as terms related to transcriptional regulation. The terms retrieved by GRNBoost2 are less specific pertaining mainly to transcriptional regulation. This indicates that, on a network level, the isoform-specific network is enriched in regulatory interactions which are relevant for the tissue in question.

**Table 2.** All enriched terms highlighted by g:Profiler for the top 100 unique target genes by weighted in-degree of the isoform-specific network computed by AlterNet and the canonical network computed by GRNBoost2 for the DCM dataset obtained using the g:Profiler R package. The terms are first ordered by ontology, then by p-value.

| Isoform-specific targets (AlterNet) |  | Canonical targets (GRNBoost2) |  |
| --- | --- | --- | --- |
| GO term | Adjusted *P*-value | GO term | Adjusted *P*-value |
| regulation of RNA metabolic process (GO:BP) | 7.66E-46 | transcription by RNA polymerase II (GO:BP) | 8.41E-48 |
| cardiocyte differentiation (GO:BP) | 5.01E-06 | chromatin (GO:CC) | 3.84E-22 |
| pattern specification process (GO:BP) | 7.57E-04 | transcription regulator complex (GO:CC) | 3.30E-13 |
| anatomical structure morphogenesis (GO:BP) | 2.36E-02 | RNA polymerase II transcription regulatory region sequence-specific DNA binding (GO:MF) | 4.91E-57 |
| atrioventricular node cell fate commitment (GO:BP) | 4.89E-02 | DNA-binding transcription factor activity (GO:MF) | 1.03E-55 |
| chromatin (GO:CC) | 2.27E-22 | transcription factor binding (GO:MF) | 8.72E-10 |
| transcription regulator complex (GO:MF) | 3.83E-08 | metal ion binding (GO:MF) | 1.01E-06 |
| DNA-binding transcription factor activity (GO:MF) | 1.02E-52 | chromatin binding (GO:MF) | 4.81E-02 |
| RNA polymerase II transcription regulatory region sequence-specific DNA binding (GO:MF) | 1.66E-52 | protein binding (GO:MF) | 8.20E-08 |
| metal ion binding (GO:MF) | 1.17E-05 |  |  |
| transcription factor binding (GO:MF) | 9.86E-04 |  |  |

Since cardiomyopathies have been linked to aberrant AS [3], we next asked if known splice factors (SFs) are enriched in the targets of the isoform-specific networks inferred for the DCM and HCM datasets in comparison to targets in the canonical GRNs computed by GRN-Boost2. Indeed, we observe a statistically significant enrichment for the DCM datasets (Figure 4). For the HCM dataset the enrichment was not significant, which could be related to the lower sample size (*n* = 28). This provides further evidence that AlterNet’s isoform-specific GRNs pinpoint to condition-specific molecular processes.

**Fig. 4.**
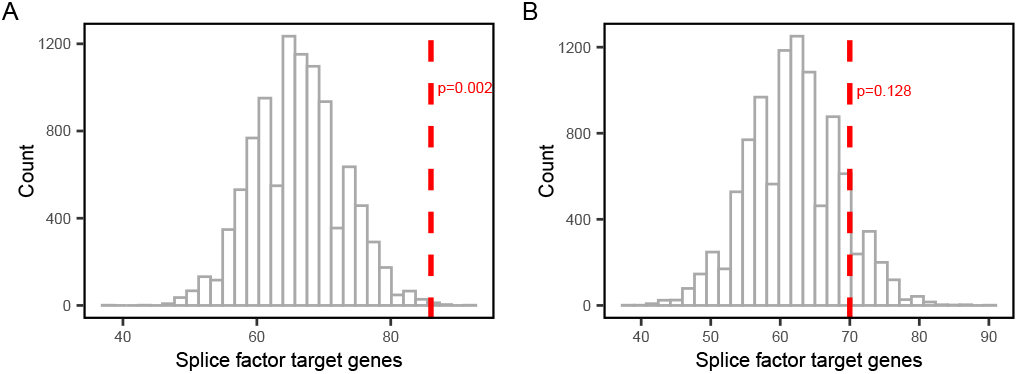
Overrepresentation analysis of splice factors in target lists of isoform-specific networks compared to canonical GRNs for the DCM (A) and HCM (B) datasets. Empirical *P*-values were computed as explained in Section 4.3.

### 2.5 Runtime

Table 3 shows AlterNet’s runtimes for the three datasets, separately for inference of the canonical and the AS-aware GRNs and all preprocessing steps. As expected, GRN inference dominates the total runtime. On average, AlterNet’s total runtimes are around two to three times higher than those of vanilla GRNBoost2, as can be seen by comparing columns “Total” and “Inference of canonical GRN” in Table 3. This shows that GRN inference with AlteNet remains feasible for realistically sized datasets.

**Table 3.** Runtimes in minutes for the three datasets.

| Dataset | Inference of canonical GRN | Inference of AS-aware GRN | Postprocessing | Total |
| --- | --- | --- | --- | --- |
| DCM | 87 | 170 | 19 | 276 |
| HCM | 103 | 123 | 42 | 268 |
| NF | 90 | 171 | 19 | 280 |

## 3 Discussion

This study addresses the limitations of conventional gene-level GRN inference by introducing the transcript-level, AS-aware pipeline AlterNet. AlterNet models TF isoforms as distinct regulators and integrates transcript-level annotations, thereby enabling finer regulatory resolution than traditional gene-level GRN inference methods. From a technical standpoint, AlterNet incorporates both canonical gene-level and isoform-level GRN inference using a variant of GRNBoost2, followed by postprocessing steps including edge classification and plausibility filtering. These steps yield a subset of high-confidence isoform-specific regulatory links that tend to be overlooked in classical GRN inference, which are then augmented with annotations such as APPRIS principal transcript status, TRIFID scores, and domain- and exon-usage labels from DIGGER. Despite these additions, runtime analysis confirmed that the computational overhead is minimal, as the majority of runtime remains associated with the inference stage itself. This demonstrates that AS-aware GRN inference is computationally scalable.

Biological validation using transcript expression data from human heart tissue — including samples from donor hearts with apparently normal function and samples from DCM and HCM patients — highlighted the biological value of transcript-level modeling. In particular, gene set enrichment on the targets of the isoform-specific interactions revealed a high number terms directly related to cardiac function, while producing a smaller number of putative regulations. This corresponds to an absolute and relative enrichment of information in the isoform-specific GRNs in comparison to canonical gene-level GRNs, making the identification of novel hypotheses less cumbersome.

Currently, only the TFs are considered at the isoform level. In the future, we intend to extend AlterNet to also model the targets of the inferred regulatory links at the level of isoforms. For this, we will incorporate SF expression into the inference process, which will allow us to jointly model the regulatory processes controlling AS and transcription. A further limitation of our study is that there are no gold standard networks which can be used to evaluate the inferred isoform-specific GRNs. Curated datasets such as CollecTRI [19] contain experimentally confirmed TF-target interactions only at the gene level and hence cannot be used to test AlterNet’s isoform-specific GRNs. The same problem applies to simulators such as scMultiSim [13] that simulate the effect of a GRN into synthetic gene expression data. In the future, we plan to investigate AS-aware GRNs on a larger scale, including on more datasets. In particular, the HCM dataset has a limited number of samples, which may lead to less granular results.

To summarize, we have shown that transcript-level GRN inference is both technically feasible and biologically informative. By leveraging isoform-level expression and functional annotation, our AlterNet pipeline uncovers regulatory relationships that remain hidden in gene-centric models. This underscores the value of incorporating AS into GRN inference to more accurately capture the complexity of transcriptional regulation in human tissues.

## 4 Methods

### 4.1 The AlterNet pipeline

#### Input and preprocessing

AlterNet requires three inputs: (i) the transcript expression matrix *X*^ℐ^ ∈ ℝ^*n×*|ℐ |^ quantified as transcripts per million (TPM), with *n* the number of samples and ℐ the set of isoforms; (ii) a mapping *σ* : ℐ → 𝒢 that assigns each isoform *i* ∈ ℐ to exactly one *g* ∈ 𝒢, where 𝒢 is the set of gene names, and (iii) a list of TFs ℛ ⊆ 𝒢 that serve as candidate regulators. To obtain the gene-level TPM matrix *X*^𝒢^ ∈ ℝ^*n×*|𝒢|^, we sum up all transcript TMPs per gene, i. e., define 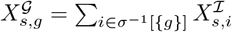 for sample *s* and gene *g*. Before constructing *X*^𝒢^, the set of transcripts is filtered such that only protein-coding transcripts are retained as columns in *X*^ℐ^. This ensures that *X*^ℐ^ and *X*^𝒢^ contain only counts from protein isoforms that can potentially become active as a TF. Moreover, we remove lowly expressed isoforms where 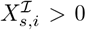 for less than *q* · *n* samples (default: *q* = 0.5), and scale the columns 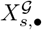 of the gene expression matrix to unit variance via *z*-score normalization to ensure that the edge weights of the inferred GRNs (see next paragraph) are comparable across different target genes.

#### GRN inference

GRN inference is performed using GRNBoost2, which has shown good performance in independent benchmarks [24]. We use a custom modification implemented in our SignifiKANTE package [31], which allows to pass the set of target genes as an additional input. Given a gene expression matrix *X* ∈ ℝ^*n×*|𝒢|^ and a set of TFs ℛ ⊆ 𝒢, GRNBoost2 formulates the GRN inference problem as a feature selection problem and creates random regression forest (RF) models *f*_*g*_ of form *X*_•,*g*_ ∼ *f*_*g*_(*X*_•,*R \*{*g*}_) to find the most predictive TFs for the expression vectors *X*_•,*g*_ of each target gene *g* ∈ 𝒢. In AlterNet, GRNBoost2 is called twice: For the first call, we set *X* = *X*^𝒢^ and thus produce a canonical gene-based GRN *G*^*c*^ = (𝒢, *E*^*c*^, *w*) with directed edges (*r, g*) ∈ *E*^*c*^ from TFs *r* ∈ ℛ ⊆ 𝒢 to target genes *g* ∈ 𝒢, where the edge weight *w*(*r, g*) corresponds to the feature importance of the TF *r* in the RF model *f*_*g*_. For the second call, we map the set of TFs ℛ to the corresponding isoforms ℛ ^ℐ^ = *σ*^−1^[ℛ] and then infer a bipartite AS-aware GRN *G*^*a*^ = (ℛ ^ℐ^ ∪𝒢, *E*^*a*^, *w*) with directed edges (*i, g*) ∈ *E*^*a*^ from TF isoforms *I* ∈ ℛ ^ℐ^ ⊆ ℐto target genes *g* ∈ 𝒢. For this, we use our customized version of GRNBoost2, which allows to train RF models *f*_*g*_ of the form 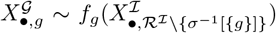. To make the pipeline more robust, both GRN inference calls are repeated *N* times (default: *N* = 10) and the frequencies and median importance weights of all edges are recorded.

#### Edge categorization

We proceed in a two step fashion to categorize the edges (*r, g*) ∈ *E*^*c*^ and (*i, g*) ∈ *E*^*a*^ in the inferred canonical and AS-aware GRNs *G*^*c*^ and *G*^*a*^. For a TF-target gene edge (*r, g*) ∈ *E*^*c*^ from the canonical GRN, let 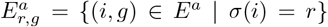 be the set of corresponding isoform-target gene edges from the AS-aware GRN (note that 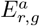 may be empty). We partition the edge set *E*^*c*^ into the set 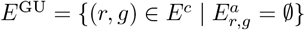 of *gene-unique* TF-target gene edges that can be found only in the canonical GRN and the set *E*^CG^ = *E*^*c*^ *\E*^GU^ of *common edges* for which a corresponding isoform-target edge exists in the AS-aware GRN. Similarly, we partition *E*^*a*^ into the sets *E*^CI^ = {(*i, g*) ∈ *E*^*a*^ | (*σ*(*i*), *g*) ∈ *E*^*c*^} and *E*^IU^ = *E*^*a*^ \ *E*^CI^ of *common* and *isoform-unique* edges (note that *E*^CG^ and *E*^CI^ can be considered gene- and isoform-level views of the same set of *common edges*). These edge sets can then optionally be filtered with importance and frequency thresholds. In the interest of obtaining stable, high-confidence results, we recommend using edges which are identified in all *N*. The common edges are further subdivided into *equivalent edges, likely isoform-unique egdes, likely gene-unique edges*, and *ambiguous edges*. Consider a pair of edges (*i, g*) ∈ *E*^CI^ and (*r, g*) ∈ *E*^CG^ with *σ*(*i*) = *r*. The two edges are categorized as equivalent if *σ*^−1^[{*r*}] = {*i*}, i. e., if *i* is the only isoform of the TF *r* present in the dataset. The edge (*i, g*) is categorized as likely isoform-unique if *w*(*i, g*)*/w*(*r, g*) ≥ *δ* and |*w*(*i, g*) −*w*(*r, g*)| ≥ Δ (the latter filter is required to remove edge pairs with generally low explainablity). The edge (*r, g*) is categorized as likely gene-unique if *w*(*i*^*′*^, *g*)*/w*(*r, g*) ≤ *δ*^−1^ and |*w*(*i*^*′*^, *g*) − *w*(*r, g*)| ≥ Δ for all 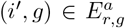. The remaining common edges are categorized as ambiguous. Δ *>* 0 and *δ >* 1 are hyperparameters that default to Δ = 1 and *δ* = 1.5. Finally, we define a derived category of *isoform-specific* edges that comprises all isoform-unique and likely isoform-unique interactions.

#### Edge filtering

Based on the edge categorization, we apply five filters that act on edges from the different categories: the *frequency filter*, the *equivalence filter*, the *dominance filter*, the *foldchange filter*, and the *importance filter*. The frequency filter requests that retained edges (*i, g*) ∈ *E*^*a*^ and (*r, g*) ∈ *E*^*c*^ should appear in at least *r* % of the *N* individual GRNs inferred by GRNBoost2. It acts on all edges independently of their category. By default, we set a very stringent threshold of *r* = 100 to focus on statistically robust edges. The equivalence and dominance filters act on isoform-unique edges (*i, g*) ∈ *E*^IU^ and gene-unique edges (*r, g*) ∈ *E*^GU^. The equivalence filter requests that, for these edges, isoforms should not be equivalent to genes. That is, edges (*r, g*) with | *σ*^−1^[{*r*}] | = 1 are removed from *E*^GU^ and edges (*i, g*) with | *σ*^−1^[{*σ*(*i*)}] | ={*i*} are removed from *E*^IU^. The dominance filter additionally removes edges (*i, g*) from *E*^IU^ where the isoform *i* accounts for at least *τ* % (default: *τ* = 90) of the genes *σ*(*i*)’s total expression and edges (*r, g*) from *E*^GU^ where *σ*^−1^[{*r*}] contains one such dominant isoform. The rationale behind these two filters is that, when there is only one or one dominant isoform for a given TF, the TF and the (dominant) isoform are (almost) statistically equivalent and the corresponding edges should hence appear in both the canonical and the AS-aware GRN. The foldchange filter acts on pairs ((*i, g*), (*σ*(*i*), *g*) ∈ *E*^CI^ × *E*^GU^ of common equivalent edges. It requests that the importance weights of the edges (*i, g*) and (*σ*(*i*), *g*) are similar and removes both (*i, g*) from *E*^CI^ and (*σ*(*i*), *g*) from *E*^GU^ if *w*(*i, g*)*/w*(*σ*(*i*), *g*) ∉ [1 − *ϵ*, 1 + *ϵ*] (default: *ϵ* = 0.5). Lastly, the importance filter requests that all edges should have a minimum importance weight to focus on highly explanatory edges. The importance threshold is calculated by computing percentiles over the relevant importance weights of all surviving edges. All edge lists are then thresholded using a common percentile threshold *ρ* (default: *ρ* = 80).

#### Functional transcript annotation

To gain a better understanding of the transcripts involved in the inferred interactions, the regulatory network inference is complemented with functional transcript annotations from the APPRIS [25] and DIGGER [14] databases. APPRIS is a database that provides annotations for splice isoforms of protein-coding genes. Isoforms are classified into categories based on multiple criteria, including evolutionary conservation, structural information, and presence of conserved functional domains. The three main categories are *principal* for isoforms with high functional evidence, *alternative* for isoforms with medium evidence, and *minor* for the remaining isoforms. Additionally, the APPRIS annotations include TRIFID scores, a learned score that assesses the functional importance of transcript isoforms, ranging from 0 (low evidence) to 1 (high evidence). It integrates evolutionary conservation, protein structure, and functional domain evidence to estimate the functional importance of individual isoforms. Finally, APPRIS provides annotations indicating if a transcript is protein-coding or not. Overall, we include the annotations of 172157 isoforms of 20445 genes. The DIGGER database provides information on protein-protein, domain-domain, and residue-level interactions for 67552 alternatively spliced isoforms of 18379 genes. We use DIGGER to retrieve full exon and Pfam domain annotations for all isoforms *i* involved in an edge (*i, g*) of the filtered AS-aware GRN, along with specific subsets of these annotations that depend on the edge categories:

– If (*i, g*) is an isoform-unique or likely isoform-unique edge, we retrieve unique exons and Pfam domains, using the APPRIS classification to determine the comparison baseline. If *i* is labeled as a minor or alternative isoform in APRIS, it is compared to the associated principal isoform *i*^*′*^ if one is available. If no principal isoform exists or if *i* itself is labeled as principal, it is compared against all other *i*^*′*^ isoforms associated with the same gene. We then annotate *i* with its exons and Pfam domains from DIGGER that do not appear in the DIGGER annotations for the baseline isoforms *i*^*′*^. The presence of unique exons may indicate distinct protein sequences and structural features, suggesting isoform-specific functional roles [1]. Similarly, unique Pfam domains suggest the presence of isoform-specific functional regions [17].
– Isoforms *i* ∈ *σ*^−1^[{*r*}] of a TF *r* for which the filtered canonical GRN contains a gene-unique or likely gene-unique edge (*r, g*) are annotated with common exons and common Pfam domains. For this, we retrieve the sets of exon and domain annotations from DIGGER for all isoforms *i* ∈ *σ*^−1^[{*r*}], and then annotate each of these isoforms with the intersections. Knowledge about the shared domains can indicate to the user why all isoforms of a gene must cooperate in order to regulate a target gene.

#### Output of the AlterNet pipeline

The final output is a set of ranked list of isoform-unique, likely isoform-unique, gene-unique, likely gene-unique, equivalent, and ambiguous regulatory interactions. Edges ranked higher in the lists are considered more plausible based on inference frequency and importance scores. Each entry includes full annotations from APPRIS and DIGGER for the source transcript, along with additional columns indicating unique or shared exon identifiers and Pfam domains. This comprehensive output facilitates downstream analysis by prioritizing regulatory interactions in which the source transcript has a higher likelihood of having an individual functional relevance. It also provides additional features that aid in the investigation of potential isoform-specific functions and resulting regulatory connections.

### 4.2 Transcript expression datasets

We used transcript expression data generated by the MAGNet consortium [5,27]. The dataset includes 366 expression profiles from the left ventricular heart tissue from non-failing donors (NF, 166 samples) and tissues harvested during surgery from patients diagnosed with heart failure. Patients diagnosed with heart failure had one of three cardiomyopathy conditions: peripartum cardiomyopathy (PPCM, 6 samples), hypertrophic cardiomyopathy (HCM, 28 samples), and dilated cardiomyopathy (DCM, 166 samples). For our study, we excluded the PPCM samples due to the limited sample availability. For all NF, HCM, and DCM samples, we performed transcript quantification on the raw data using Kallisto [4], thus obtaining TPMs for all samples and transcripts. Kallisto is a convenient tool for this application, as it allows the summation of TPMs to obtain accurate total abundance measures [28]. The unfiltered datasets contain values for 57773 genes and 188753 transcripts. We then removed all isoforms which are expressed in less than 50% of the samples, independently for the DCM, HCM, and NF datasets. The resulting dataset statistics are shown in Table 4.

**Table 4.** Numbers of samples, genes, TFs, and TF isoforms in the three datasets used in this study after removal of isoforms expressed in less than half of the samples of the datasets. TFs are a subset of the genes.

| Dataset | Samples | Genes | TFs | TF isoforms |
| --- | --- | --- | --- | --- |
| DCM | 166 | 15955 | 1539 | 4191 |
| HCM | 28 | 16116 | 1549 | 4411 |
| NF | 166 | 15827 | 1530 | 4332 |

### 4.3 Evaluation strategies

#### Computation of empirical P-values based on unique exons and Pfam domains

Let *X*^ℐ^ be one of the three transcript expression datasets, *E*^⋆^ ⊂ *E*^*a*^ be the subset of isoform-specific edges obtained for *X*^ℐ^ after applying all filters, and ℐ_*E*_⋆ = {*i* ∈ ℐ | ∃*g* ∈ 𝒢: (*i, g*) ∈ *E*^⋆^}be the set of involved TF isoforms. Using the domain annotations from DIGGER and the isoform categories from APPRIS, we determined the number *u*(ℐ_*E*_⋆) = |{*i* ∈ ℐ_*E*_⋆ | *i* is not principal and has unique domain}| of non-principal isoforms *i* ∈ ℐ_*E*_⋆ that have a domain not present in the principal isoform, and the number *m*(ℐ_*E*_⋆) of non-principal isoforms *i* ∈ ℐ_*E*_⋆ where a domain from the principal isoform is missing. Then, we randomly sampled 1000 size-matched edge sets 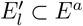 with 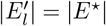 from the unfiltered AS-aware GRN, and analogously computed background distributions of 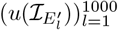 and 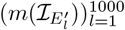 of unique exon and domain counts. Based on those, we computed empirical *P*-values

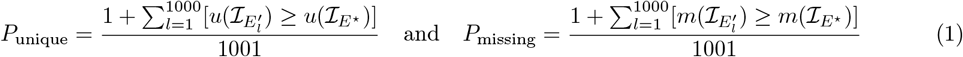

that are small if the domain annotations from DIGGER yield evidence that non-principal isoforms that appear as regulators in the filtered isoform-specific networks have functional profiles that differ from the functional profiles of the corresponding principal isoforms. Analysis of SF overrepresentation in the isoform-specific networks in comparison to the canonical GRNs was carried out similarly: We first compiled a set of 233 SF from SpliceAid-F [7], SplicingLore [22], [29], and the KEGG splicesome pathway and removed all auto-regulatory edges from the networks to avoid biasing the target genes, as TFs can also be targets. Then, analogously to the analysis of unique and missing domains, we compared the number of SFs among the targets of the isoform-specific GRNs to empirical distributions of SF counts in 1000 size-matched randomly sampled subsets of targets in the canonical GRNs.

#### Functional enrichment analysis

We used two variants of functional enrichment analysis to assess the functional plausibility and relevance of the set *E*^⋆^ of isoform-unique or likely isoform-unique interactions. For the first variant, we used the annotations from DIGGER to construct the set 𝒟 of Pfam domains associated with any of the isoforms *i* ∈ ℐ present in the dataset, as well as the subset 𝒟^⋆^ ⊂ 𝒟 of domains that are associated exclusively with isoforms *i* ∈ ℐ_*E*_⋆ that act as regulator in the filtered set of isoform-specific interactions. Moreover, for each domain *d* ∈ 𝒟, we retrieved the set GO(*d*) of associated GO terms. For each GO term *t* ∈ ∪_*d*∈*D*_ GO(*d*), we then carried out a one-sided hypergeometric test to assess if *t* is overrepresented in 𝒟^⋆^, given its overall frequency in 𝒟. Subsequently, we adjusted the resulting *P*-values for multiple testing using Bonferroni correction. For the second variant, we extracted the top 200 target genes (sorted by weighted in-degree) from both *E*^⋆^ and the canonical GRN *G*^*c*^. For both sets of target genes, we then carried out gene set enrichment analysis against the GO database using the g:Profiler R package [10], and reported the obtained multiple testing-corrected *P*-values.

#### Runtime evaluation

To evaluate the computational performance of the developed inference and annotation pipeline, we carried out a runtime evaluation. All experiments were executed on a shared workstation equipped with a dual-socket AMD EPYC 7402 24-core processor with 96 threads and 528 GB RAM. Using Python’s `time` module, we separately recorded the runtime for the GRN inferrence step and the filtering pipeline.

## 5 Code and data availability

AlterNet is available at https://github.com/bionetslab/AlterNet, scripts to reproduce the reported results at https://github.com/bionetslab/alternet-manuscript. MAGNet data was obtained from NCBI Gene Expression Omnibus (https://www.ncbi.nlm.nih.gov/geo/query/acc.cgi?acc=GSE141910). The APPRIS database was downloaded from https://appris.bioinfo.cnio.es/#/downloads and the DIGGER database from https://exbio.wzw.tum.de/digger/download/. A mapping between Ensembl gene IDs and gene names was obtained using BioMart [9]. A list of human TFs was obtained from https://resources.aertslab.org/cistarget/tf_lists/. Pfam domain clades were obtained from http://ftp.cbi.pku.edu.cn/pub/databases/Pfam/latest_release/. The domain to GO term mapping was obtained from https://current.geneontology.org/ontology/external2go/pfam2go. The KEGG splicesome was obtained from https://www.kegg.jp/pathway/ko03040.

## Acknowledgments

D. B. B. was funded by the German Research Foundation (DFG, 516188180). D. B. B. and A. H. were funded by the German Federal Ministry of Research, Technology and Space (BMFTR, 031L0309A). We thank anonymous reviewers for their comments, we are currently preparing an extension of the pipeline based on their comments.

## Author contributions statement

J. H. implemented the analysis pipeline and analyzed the data. J. W. implemented the modified version of GRNBoost2. Z. D. compiled the list of splice factors. O. T. provided feedback on the pipeline and preprocessed MAGNet data. A. H. conceptualized the study and implemented the analyses. J. H., A. H., and D. B. B. drafted the manuscript. All authors read and reviewed the manuscript.

## Disclosure of Interests

All authors declare no competing interests.

## Notes

https://github.com/bionetslab/alternet

### Competing Interest Statement

The authors have declared no competing interest.

### Summary of Updates

Text corrections to improve clarity including spelling errors in several sections.

