## Supplementary material for "AlterNet: Alternative splicing-aware gene regulatory network inference": AlterNet_supplement (from another version of this preprint: version 1, 2025-11-22)

### Appendix

Juliane Hoffmann[1], Julia Wallnig[1], Ziheng Dai[1], Olga Tsoy[2], David B. Blumenthal[1], and Anne Hartebrodt[1]

[1] Biomedical Network Science Lab, Department Artificial Intelligence in Biomedical Engineering, Friedrich-Alexander-Universität Erlangen-Nürnberg, Nürnberger Str. 74, 91052 Erlangen, Germany
{juliane.hoffmann,julia.wallnig,ziheng.dai,david.b.blumenthal,anne.hartebrodt}@fau.de
[2] Department of Computer Science, Vrije Universiteit Amsterdam, De Boelelaan 1111, 1081 HV, Amsterdam, The Netherlands


### A Supplementary tables and figures

Table A.1: **HCM.** Top 10 enriched terms highlighted by g:Profiler for the top 100 unique target genes by weighted in-degree of the isoform-specific network computed by AlterNet and the canonical network computed by GRNBoost2 for the HCM dataset obtained using the g:Profiler R package.

| **Isoform-specific targets (AlterNet)** | | **Canonical targets (GRNBoost2)** | |
|---|---|---|---|
| **GO term** | **Adjusted $P$-value** | **GO term** | **Adjusted $P$-value** |
| regulation of DNA-templated transcription | 0.00518 | transcription by RNA polymerase II | 8.41E-48 |
| regulation of primary metabolic process | 0.0184 | chromatin | 3.84E-22 |
| morphogenesis of an epithelial sheet | 0.0248 | transcription regulator complex | 3.3E-13 |
| regulation of intracellular steroid hormone receptor signaling pathway | 0.0414 | RNA polymerase II transcription regulatory region sequence-specific DNA binding | 4.91E-57 |
| transcription factor binding | 0.000986 | DNA-binding transcription factor activity | 1.03E-55 |
| sequence-specific double-stranded DNA binding | 0.0122 | transcription factor binding | 8.72E-10 |
| transcription regulator activity | 0.0144 | metal ion binding | 0.00000101 |
| | | chromatin binding | 0.0481 |

Table A.2: **NF.** Top 10 enriched terms highlighted by g:Profiler for the top 100 unique target genes by weighted in-degree of the isoform-specific network computed by AlterNet and the canonical network computed by GRNBoost2 for the NF dataset obtained using the g:Profiler R package.

| **Isoform-specific targets (AlterNet)** | | **Canonical targets (GRNBoost2)** | |
|---|---|---|---|
| **GO term** | **Adjusted $P$-value** | **GO term** | **Adjusted $P$-value** |
| regulation of transcription by RNA polymerase II | 2E-40 | transcription by RNA polymerase II | 8.41E-48 |
| animal organ development | 0.0000907 | chromatin | 3.84E-22 |
| His-Purkinje system cell differentiation | 0.000862 | transcription regulator complex | 3.3E-13 |
| organ growth | 0.025 | RNA polymerase II transcription regulatory region sequence-specific DNA binding | 4.91E-57 |
| pattern specification process | 0.0285 | DNA-binding transcription factor activity | 1.03E-55 |
| retinoic acid receptor signaling pathway | 0.0454 | transcription factor binding | 8.72E-10 |
| atrioventricular node cell fate commitment | 0.0479 | metal ion binding | 0.00000101 |
| nucleus | 2.68E-20 | chromatin binding | 0.0481 |
| transcription regulator complex | 4.59E-11 | | |
| DNA-binding transcription factor activity | 8.41E-50 | | |
| RNA polymerase II transcription regulatory region sequence-specific DNA binding | 1.37E-49 | | |
| metal ion binding | 0.000000019 | | |
| transcription factor binding | 0.0000038 | | |

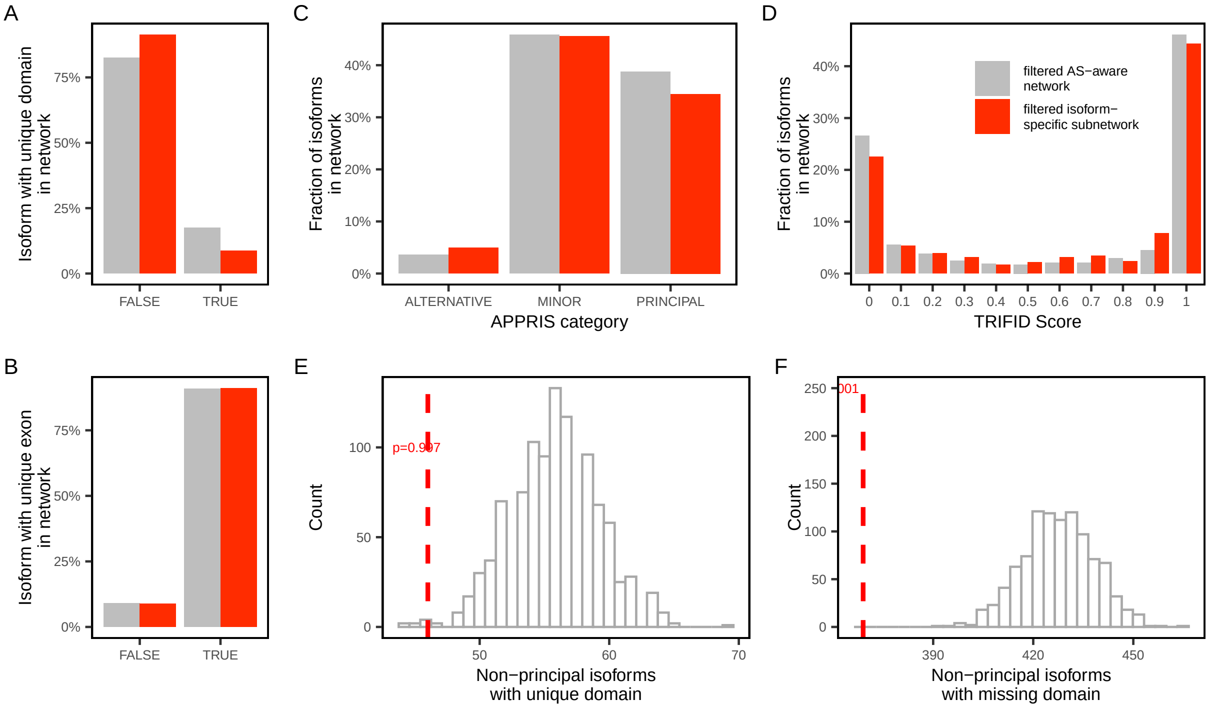


Fig. A.1: **HCM.** Analysis of TF isoforms in AlterNet's filtered isoform-specific network obtained for the DCM dataset in comparison to the filtered AS-aware network. (A, B) Numbers of isoforms with unique exons and unique domains. (C, D) Distributions of the APPRIS isoform categories TRIFID scores. (E, F) Numbers of non-principal isoforms in the isoform-specific network with unique domains (E) and missing domains (F) against background distributions generated by randomly sampling an identical number of edges from the filtered AS-aware GRN. Empirical $P$-values were computed as explained in section 4.3. in the main text.

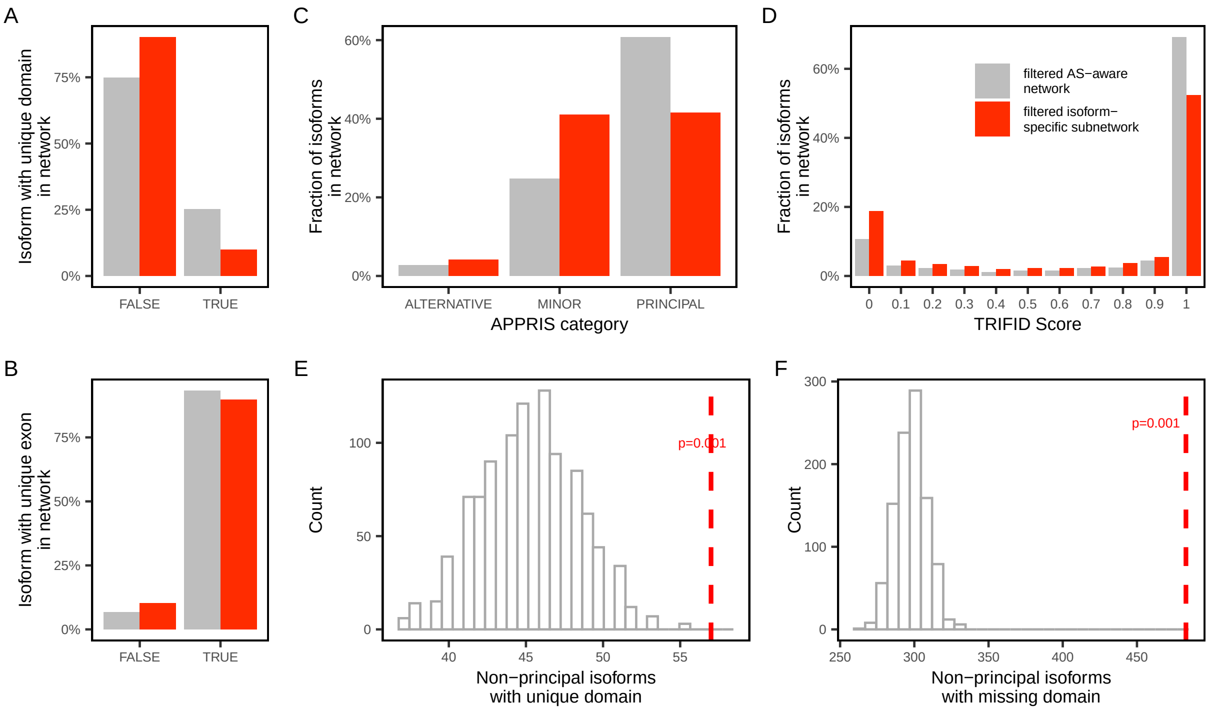


Fig. A.2: **NF.** Analysis of TF isoforms in AlterNet's filtered isoform-specific network obtained for the DCM dataset in comparison to the filtered AS-aware network. (A, B) Numbers of isoforms with unique exons and unique domains. (C, D) Distributions of the APPRIS isoform categories TRIFID scores. (E, F) Numbers of non-principal isoforms in the isoform-specific network with unique domains (E) and missing domains (F) against background distributions generated by randomly sampling an identical number of edges from the filtered AS-aware GRN. Empirical $P$-values were computed as explained in 4.3 in the main text
